# Widespread coding of navigational variables in prefrontal cortex

**DOI:** 10.1101/2022.10.13.512139

**Authors:** David J-N. Maisson, Benjamin Voloh, Roberto Lopez Cervera, Indirah Conover, Mrunal Zambre, Jan Zimmermann, Benjamin Y. Hayden

**Affiliations:** Department of Neuroscience, Center for Magnetic Resonance Research, Center for Neuroengineering, Department of Biomedical Engineering, University of Minnesota, Minneapolis MN 55455

**Keywords:** navigation, spatial selectivity, head direction tuning, mixed selectivity, prefrontal cortex

## Abstract

To navigate, we must represent information about our place in the environment. Traditional research highlights the role of the hippocampal complex in this process. Spurred by recent research highlighting the widespread cortical encoding of cognitive and motor variables previously thought to have localized function, we hypothesized that navigational variables would be likewise encoded widely, especially in the prefrontal cortex, which is often associated with control of volitional behavior. We recorded neural activity from six prefrontal structures while macaques performed a foraging task in an open enclosure. In all six regions, we found strong encoding of allocentric position, head direction, egocentric boundary distance, and linear and angular velocity. These encodings were not accounted for by distance or time to reward. Strength of coding of all variables increase along a ventral-to-dorsal gradient. Together these results argue that encoding of navigational variables is not localized to the hippocampal complex and support the hypothesis that navigation is continuous with other forms of flexible cognition in the service of action.

## INTRODUCTION

To move successfully in the world, mobile organisms must represent where they are, where they are going, and where important features in the world are. In other words, they must navigate. The majority of neuroscientific research highlights the role of the hippocampus and adjacent structures in navigation (Hartley et al., 2014; Rolls and Wirth, 2018; Moser et al., 2008; McNaughton et al., 2006), including in more general abstract mapping (Stachenfeld et al., 2017; Schuck and Niv, 2019). This work supports what is essentially a modular view of navigation, that is, it supports the view that navigation results from information computed in a dedicated set of anatomically circumscribed circuits (Epstein et al., 2017; Poulter et al., 2018). An alternative view would be that navigation relies on a suite of more general cognitive abilities, including generalized mapping functions, such as the encoding of task structure and latent environmental variables.

The modular approach to understanding functional neuroanatomy has been challenged by a growing set of studies that highlight the broad distribution of variables otherwise thought to be more narrowly circumscribed. These include motor signals (Musall et al., 2019a and b; Stringer et al., 2019; Steinmetz et al., 2019) and reward signals (Vickery et al., 2011; Shin et al., 2021; Ottenheimer et al., 2022). These findings raise the possibility that like other cognitive functions, navigation may also be more distributed than previously assumed, perhaps as part of a gradient characterized by gradual untangling rather than strict functional borders (Fine et al., 2022; Fuster, 2000 and 2001; Yoo et al., 2018). Thus, evidence of widespread distribution of navigational codes would support their posited role in anchoring elements of cognition to maps of embodied space. In this view, navigation is a special case of the more general problem of representing linkages between variables (Behrens et al., 2018; Epstein et al., 2017; Whittington et al., 2020; Stachenfeld et al., 2017; Schuck and Niv, 2019).

A small but growing body of research suggests that, at least in rodents, there may be navigational functions outside of the hippocampal complex, including within the prefrontal cortex or its homologues (Patai & Spiers, 2021; Maisson et al., 2022). For example, neurons in somatosensory cortex, orbitofrontal cortex, and piriform cortex encode current and future spatial positions as well as the location of targets in the environment (Basu et al., 2021; Long and Zhang, 2021; Poo et al., 2022; Wikenheiser et al., 2021). This work raises the possibility that navigational functions may likewise be widespread across the prefrontal cortex. However, this work is far from definitive, in part because of the distinct navigational strategies and preferred sensory modalities in rodents owing to a highly specialized navigational system, which may not generalize to primate. Moreover, the extent to which rodent prefrontal areas serve as functional homologies to those of primates remains unclear (Heilbronner et al., 2016).

Theories about the function of the prefrontal cortex seldom involve a frank navigational role. Instead, they typically involve functions that may only indirectly support navigation, including control of action, planning, resolving conflict, and comparing values (Fuster, 2000 and 2001; Miller & Cohen, 2001; Knight and Stuss, 2002). Often, however, these areas support more abstract mapping functions that augment space by anchoring to it an executive control-type function. For example, the roles for space typically assigned to PFC include the encoding of goal location, navigational action planning and landmark vector representations, and spatial working memory (Martinet et al., 2011; Strait et al., 2016; Yoo et al., 2018; Bicanski & Burgess, 2020; and as reviewed both in Ikkai & Curtis, 2011 and in Behrens et al., 2018). Yet, the involvement of PFC structures in pure navigational encoding, as well as whether these codes are distributed widely across PFC structures, remains unclear.

We developed a novel naturalistic paradigm for recording neural and behavioral data from freely navigating rhesus macaques (Bala et al., 2020). As rhesus macaques performed a freely moving foraging task, we recorded neuronal activity in six brain areas: orbitofrontal cortex (OFC), dorsal anterior cingulate cortex (dACC), supplementary motor area (SMA), ventrolateral prefrontal cortex (vlPFC), dorsolateral prefrontal cortex (dlPFC), and dorsal premotor cortex (PMd). In all six structures, we found selectivity for key navigational variables: allocentric position, head orientation, egocentric boundary distance, and angular and linear velocity. Proportions of neurons responding were roughly the same as those encoding more traditional cognitive variables, such as reward number and feeder identity. In all areas, neurons encoding navigational variables did not form functionally specialized subpopulations. Finally, we found that distributed encoding of both navigational and non-navigational task variables followed functional gradients in which stronger coding was found in more dorsal areas. Together these results support the idea that navigational functions are supported by widespread activity of neurons, including those in several prefrontal structures.

## RESULTS

### Behavioral and neural recordings

Rhesus macaque subjects performed a novel *freely-moving foraging task* in a large (245 × 245 × 275 cm) enclosure (**Figure 1A, Methods**). The enclosure contained four small barrels (height: 78.8 cm; diameter: 46.5 cm) located in the corners and was otherwise open. The enclosure contained reward stations (typically four, but on a few occasions two or three; see **Methods**). These reward stations contained touchscreens, juice reservoirs, dispenser tubes, and levers (**Figure 1B**). They were positioned at fixed locations of varying heights. Otherwise, subject behavior was physically unconstrained within this large environment.

**Figure 1.**
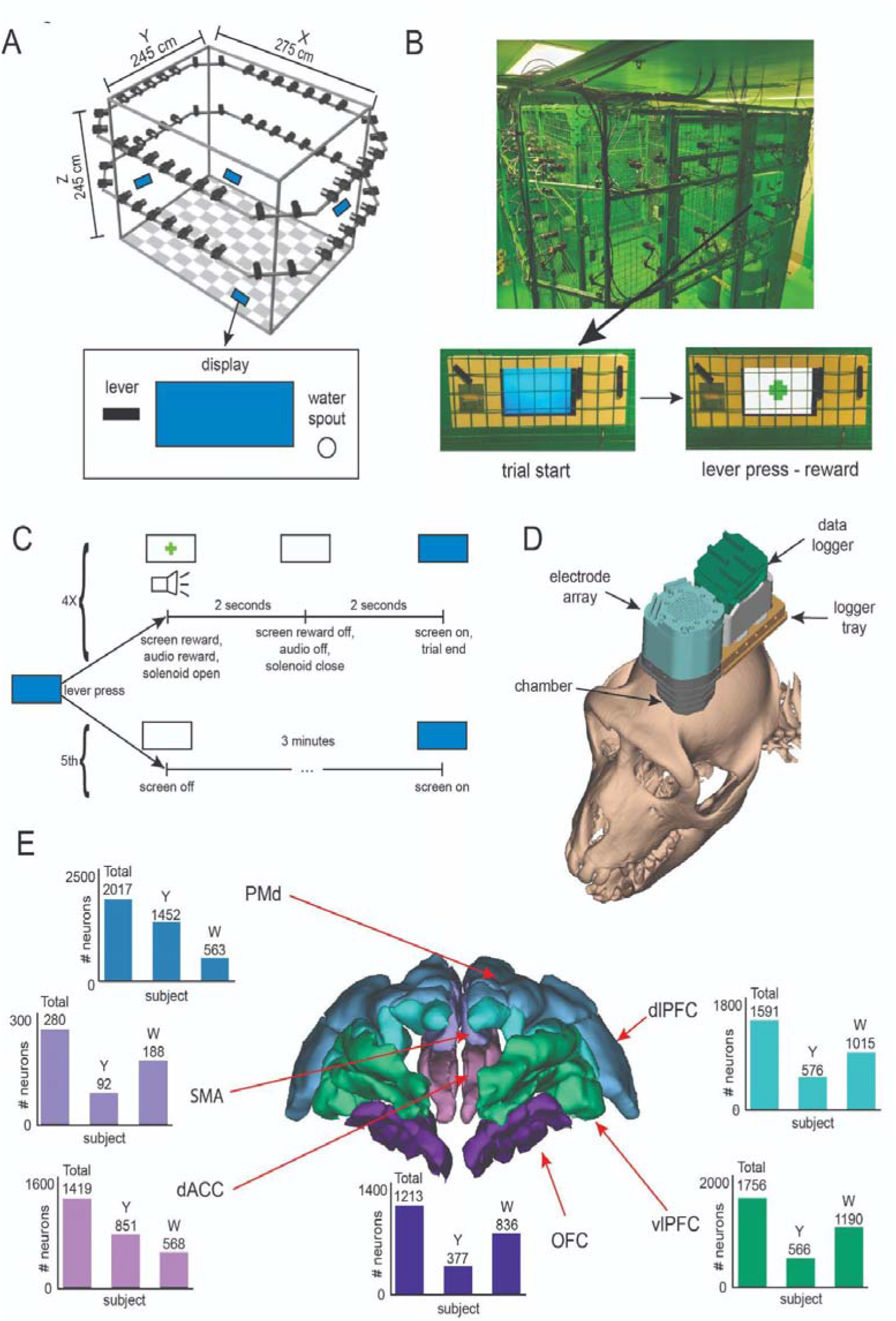
Behavior and neural recordings. **(A)** Schematic of the arena. Checkerboard patterns indicate floor. Black rectangles denote cameras. Blue rectangles indicate the approximate locations of the feeding stations. The lower panel shows a cartoon depiction of the feeding station, with the display monitor at the center showing solid blue. **(B)** Photograph of the arena. **(C)** Schematic of the available, novel *freely-moving foraging task*. **(D)** 3D model of the recording system superimposed on a subject’s cranium. **(E)** 3D rendering of the prefrontal areas from which neural data were recorded. Bars: number of neurons recorded from each region.

The computer displays were set to a solid blue screen at the start of each daily session. If the subject approached a feeding station (hereafter, a “patch”) and pressed the lever, the juice tube first provided an immediate aliquot of preferred liquid reward (typically water, always 1.5 mL). At that time, the display changed from a solid blue screen to a white screen with a solid green plus symbol in the center (**Figure 1C**). A steady (two second) tone played. Next, the tone stopped and the screen turned solid white. Following a 2-second interval, the screen returned to solid blue to indicate that the lever had been reactivated. The subject could repeat this process. After the fifth lever-press in a row at a given patch, the patch deactivated for 3 minutes. The deactivation was signaled by the patch immediately displaying solid white, with no auditory or visual cue.

We recorded the behavior of our subjects using a recently developed camera-based tracking system (*OpenMonkeyStudio*, Bala et al., 2020). This system provides estimates of the positions of 13 body landmarks in every frame of video in three dimensions. We recorded neural activity using a locally mounted data logger attached to a multi-electrode (n=128 electrodes) array with independently moveable electrodes (**Figure 1D, Methods**).

We recorded a total of 9,920 neurons over 196 sessions. Of these, 1,644 neurons were excluded from the following analyses because preliminary investigations indicated that their sessions had tracking that was too noisy for our purposes (see **Methods** for exclusion criteria). The remaining 8,276 neurons were recorded from six structures in the prefrontal cortex: OFC, dACC, SMA, vlPFC, dlPFC, and PMd (**Figure 1E**).

### Activity of prefrontal neurons can be fitted by a linear-nonlinear encoding model

We examined the selectivity of neuronal firing to a suite of navigational and task variables. To do this, we adapted a linear-nonlinear Poisson-distributed generalized additive model (LN-GAM) to estimate tuning functions simultaneously for 9 distinct variables (method was developed by Hardcastle et al., 2017 and used in Laurens et al., 2019; Vinepinsky et al., 2020; Mao et al., 2021; Yoo et al., 2020 and 2021). In our case, these variables are (1) lever pressing (vs. not pressing, which was equivalent to reward), (2) 2D position of the subject’s body on the ground plane of the enclosure, (3) head elevation, (4) absolute (compass-wise) head direction, (5) head tilt, (6) egocentric distance to enclosure boundaries, (7) egocentric distance to feeder locations, (8) angular velocity (the speed of change in the subjects’ compass-wise orientation), and (9) linear velocity of the subject’s body. These variables are the same used in previous of navigational tuning in the macaque hippocampus, and defined the same way (Laurens et al., 2019; Mao et al., 2021).

The process for identifying tuning simultaneously accounts for tuning to complex combinations of variables and tuning stability across the session (see **Methods** for greater detail). Briefly, the activity of each neuron is fitted to each variable, and each combination of variables, and compared both to the previous combination and to a null distribution to determine significance (see **Methods** and Hardcastle et al., 2017). These fits are cross-validated both within time blocks and across all time blocks for the full session. The process for determining the best-fitting model penalizes over-fitting and favors the one with the simplest possible combination of variables. To characterize encoding of basic navigational variables, we considered the prevalence of tuning to each variable in turn across neurons in each structure.

### Neurons in all six regions strongly encode egocentric position

Subjects tended to make use of a fairly large proportion of the available space in the cage. To quantify occupancy, we segmented the surface area (top-down view of the X and Y axis) of the arena into ~50 cm^2^ bins (n = 36 bins). For each bin, during each session, we computed the amount of time during which the subject’s position (specifically, its head position) was within each bin. On average across all sessions, subjects visited the entire surface area of the arena, spending an average of 2.78 ± 0.55% of the session in each bin (**Figure 2A**). We also binned the elevation (Z axis) and found that subjects visited the full range of the cage height, spending an average of 4.17 ± 0.91% of the session at each elevation, though they tended to favor an elevation between 100-200 cm (**Figure 2A**), corresponding to the height of the barrels. It is worth noting that much of the volume of the arena cannot be occupied for extended periods because of the limited number of platforms and climbing surfaces.

**Figure 2.**
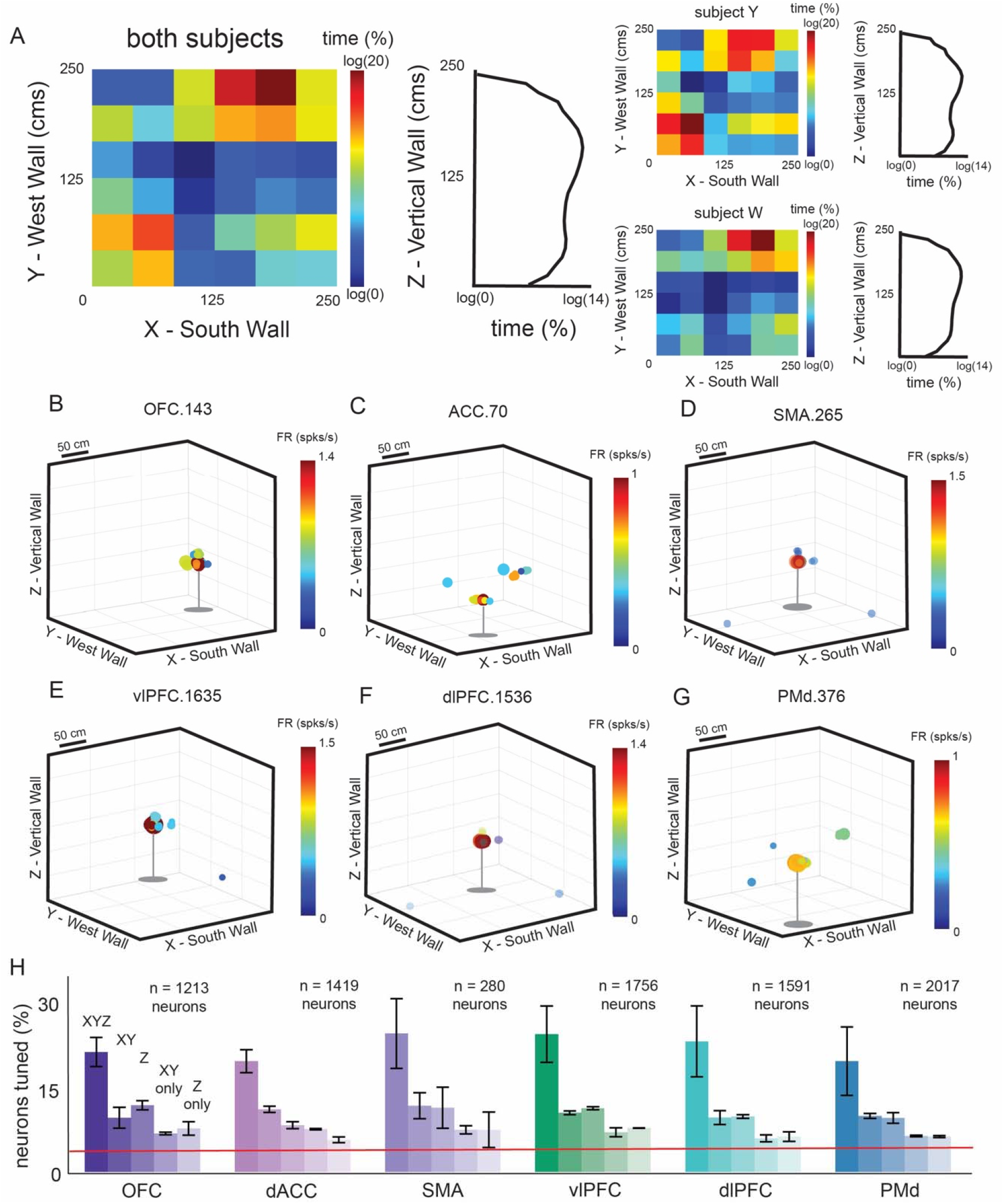
Allocentric position encoding. **(A)** Occupancy heat maps depicting the average proportion of the session spent in each 50 cm^2^ bin. X: E-W axis of the arena; Y: N-S axis. Color: proportion of the session time in each bin; right: occupancy plot for elevation axis. **(B-G)** Rate maps of a sample neuron from each structure that was determined to be significantly tuned to 3D position (2D position and tracked height, simultaneously). Color: neural activity, computed as the occupancy-normalized firing rate (spikes/second). **(H)** The proportion of total recorded neurons in each structure (collapsed across subjects) tuned to 3D position (XYZ), 2D position (XY only), tracked height (Z only), single selectivity to 2D position alone, and single selectivity to height alone. Red line: chance. Bars indicate standard error of the mean.

We examined the selectivity of neuronal firing rates to the position of the subject in the enclosure (*position encoding*). To do this, we identified neurons whose tuning profiles, in the fully fitted LN-GAM, included selectivity to position variables. In keeping with a previous study of spatial tuning in the macaque hippocampus, we defined neuronal tuning to *3D position* as the simultaneous selectivity to both the head’s *2D position* (X and Y axes, **Figure 1A**) and *head elevation* (Z axis; Mao et al., 2021).

We found that individual neurons in all six regions of the prefrontal cortex robustly encode the position of the subject in space. For example, neuron *OFC.11* showed a low baseline firing rate (0.13 spikes/sec) but showed reliably increased firing rates (1.8 spikes/sec) when the subject entered the northeast, upper corner of the enclosure (**Figure 2B**). Similarly, neuron *ACC.315* showed increased firing rates when the subject entered toward the southeast corner, elevated roughly half the height of the enclosure (**Figure 2C**). Responses of six example neurons are shown in **Figure 2D-G**.

Overall, we found that spatial selectivity is quite common in these prefrontal regions. For example, in OFC, 42.87% of neurons showed some selectivity to spatial position (Y: 50.13%; W: 39.59%). Specifically, 21.35% (n = 259/1213) of neurons carry information about *3D position*, defined as simultaneous selectivity to *2D positio*n and *head elevation* (this proportion is significantly greater than chance; *p* < 0.0001, binomial test). This pattern was observed in both subjects individually (Y: 24.93%; W: 19.74%). Beyond these 3D-tuned neurons, an additional 9.65% of OFC neurons were tuned to *2D position*, but not *head elevation* (Y: 12.22%; W: 8.49%). These proportions themselves are also significant (binomial test, *p* < 0.0001 in all cases). Similarly, an additional 11.87% of OFC neurons were tuned to *head elevation*, but not *2D position* (Y: 12.99%; W: 11.36%). We found similar results across all 5 other structures; most notably, a statistically significant proportion of neurons showed selectivity for *3D position* in all regions (**Figure 2H**). As a confirmation of tuning stability, we computed Pearson’s correlation between encoding magnitudes for each neuron to a given variable during the first and second half of the session. We found that, in all 6 structures, position tuning in the first half of the session was positively correlated with that in the second half of the session (OFC: r^2^ = 0.903; ACC: r^2^ = 0.865; SMA: r^2^ = 0.504; vlPFC: r^2^ = 0.757; dlPFC: r^2^ = 0.608; PMd: r^2^ = 0.563; *p* < 0.0001, in all cases). This finding indicates that, at least within a session, there is continuity in representational frame for these neurons.

### Neurons in all six regions encode allocentric head direction

One hallmark of navigational encoding is the tuning of neural activity to allocentric head orientation, also commonly called head direction tuning (Taube et al., 1990). Head direction tuning has been observed in structures of the broader hippocampal formation as well as thalamic nuclei known to interact with the hippocampus (as reviewed in Taube et al., 1998), and is observed in macaque hippocampus as well (Mao et al., 2021). To assess head direction selectivity in our six prefrontal structures, we used our tracking data to determine the allocentric roll, pitch, and yaw angles of the head in volumetric space at every moment (see **Methods**; as above, we used the same definitions for these terms as Mao et al., 2021).

We first confirmed that subjects head positions sampled the environment. To do this, we segmented the allocentric yaw angles into 6-degree bins (n = 60 bins, from 0-degrees to 360-degrees, 0-degrees representing due East). For each bin, during each session, we computed the amount of time during which the subject’s *head direction* (specifically, the allocentric yaw angle from a top-down view of the arena) was within each bin (**Figure 3A**). We also segmented the allocentric pitch angles into 6-degree bins (n = 30 bins, from 0-degrees to 180-degrees, 90-degrees representing an orientation directly forward). Subjects favored (an average of 63.61 ± 0.59% of the session) orienting their heads forward at an angle between 72 and 132 degrees (**Figure 3A**). For the roll angle, 90-degree represents an orientation of the head being straight up and down. Subjects overwhelmingly favored (an average of 69.51 ± 0.09% of the session) orienting their heads at an angle between 60 and 120 degrees (**Figure 3A**).

**Figure 3.**
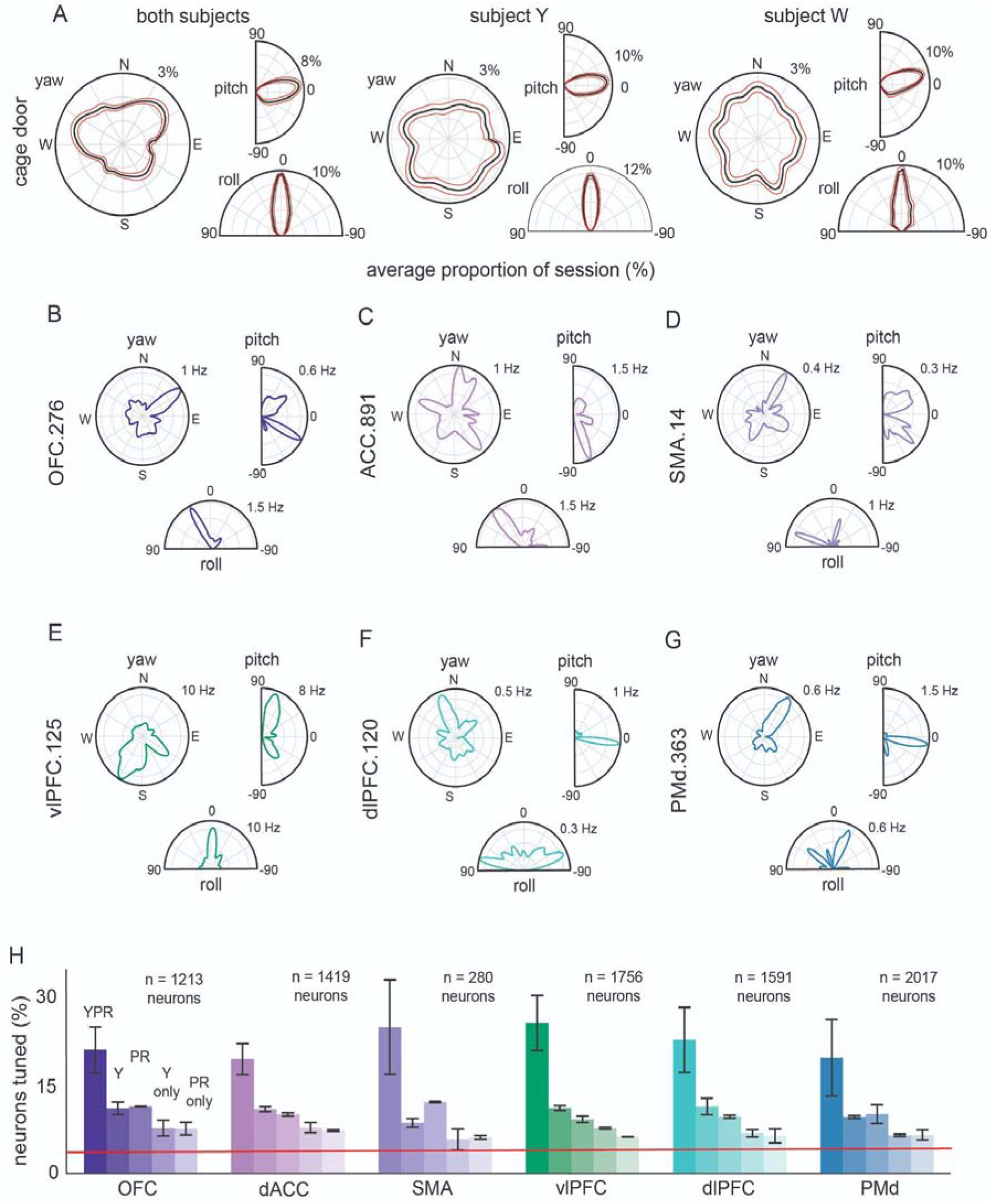
Allocentric head orientation coding in volumetric space. **(A)** Occupancy histograms depicting the distribution across sessions of the proportion of the total orientations recorded. R: proportion orientation bins occupied at least once during a session; theta: orientation. **(B-G)** Polar rate maps of a sample neuron with significant tuning. R: occupancy-normalized firing rate (spikes/second). **(H)** Proportion of total recorded neurons in each structure (collapsed across subjects) tuned to 3D orientation (yaw, pitch and roll), head direction (yaw only), head tilt (pitch and roll), single selectivity to head direction, and single selectivity to head tilt. Red line: chance.

We defined neuronal tuning to *3D orientation* as the selectivity to both the azimuthal *head direction* (yaw axis from a top-down view of the arena) and *head tilt* (both pitch and roll angles from a horizontal view of the arena; the same definition was used in Mao et al., 2021). We found that individual neurons in the prefrontal cortex encode the allocentric orientation of the subject in space. For example, neuron *OFC.276* showed a low baseline firing rate (0.04 spikes/sec) but showed reliably increased firing rates (roughly 1.03 spikes/sec) when the subject’s head was oriented toward the northeast corner of the enclosure (*head direction*), while the head was tilted slightly down and to the left (*head tilt*). (**Figure 3B**). Likewise, neuron *vlPFC.125* showed increased firing rates when the subject faced towards the southwest corner of the enclosure, with their head tilted up and straight (**Figure 3C**). Responses of six example neurons are shown in **Figure 3D-G**.

Overall, we found that allocentric orientation selectivity was quite common in these prefrontal regions. For example, in OFC, 43.53% of neurons showed some selectivity to *3D orientation* (Y: 47.48%; W: 41.75%). Specifically, 21.12% (n = 256/1213) of neurons carry information about *3D orientation*, defined as simultaneous selectivity to *head direction* and *head tilt* (this proportion is greater than chance at 5%; *p* < 0.0001, binomial test). This pattern was observed in both subjects individually (Y: 26.52%; W: 18.66%). An additional 11.05% of OFC neurons were tuned to *head direction*, but not *head tilt* (Y: 9.55%; W: 11.72%). Similarly, an additional 11.38% of OFC neurons were tuned to *head tilt*, but not *head direction* (Y: 11.41%; W: 11.36%). We found similar results across all 5 other structures; specifically, a statistically significant proportion of neurons showed selectivity for *3D orientation* in all regions (**Figure 3H**). As a confirmation of stability in tuning, we computed correlation between encoding magnitudes for each neuron to a given variable during the first and second half of the session. We found that, in all 6 structures, orientation tuning in the first half of the session was positively correlated with that in the second half of the session (OFC: r^2^ = 0.922; ACC: r^2^ = 0.846; SMA: r^2^ = 0.504; vlPFC: r^2^ = 0.774; dlPFC: r^2^ = 0.624; PMd: r^2^ = 0.548; *p* < 0.0001, in all cases).

### Encoding of boundary distance, landmark distance, and linear and angular velocity

We tested four other variables using our LN-GAM: egocentric boundary distance, egocentric landmark distance, linear velocity, and angular velocity (as above, these variables were also tested in Mao et al., 2021). Tuning to these variables was, again, defined by identifying which neurons had fully fitted models that included these variables in their selectivity profiles. We found coding of all four of these variables in all six of our regions.

Egocentric boundary distance is a subject’s current distance to the nearest boundary of the environment (Mao et al., 2021). In keeping with this approach, we defined egocentric boundary distance in polar coordinates on the horizontal plane. Briefly, we computed the radial distance between the subject’s head and the nearest wall, and the angle relative to the center of the room (see **Methods**). We found that a significant proportion of neurons in OFC were tuned to egocentric boundary distance (32.15%; *p* < 0.0001; binomial test). We found similar results in all 5 other areas (**Figure 4**). We found that, in all 6 structures, egocentric boundary tuning in the first half of the session was positively correlated with that in the second half of the session (OFC: r = 0.94; ACC: r = 0.93; SMA: r = 0.73; vlPFC: r = 0.86; dlPFC: r = 0.78; PMd: r = 0.74; *p* < 0.0001, in all cases).

**Figure 4.**
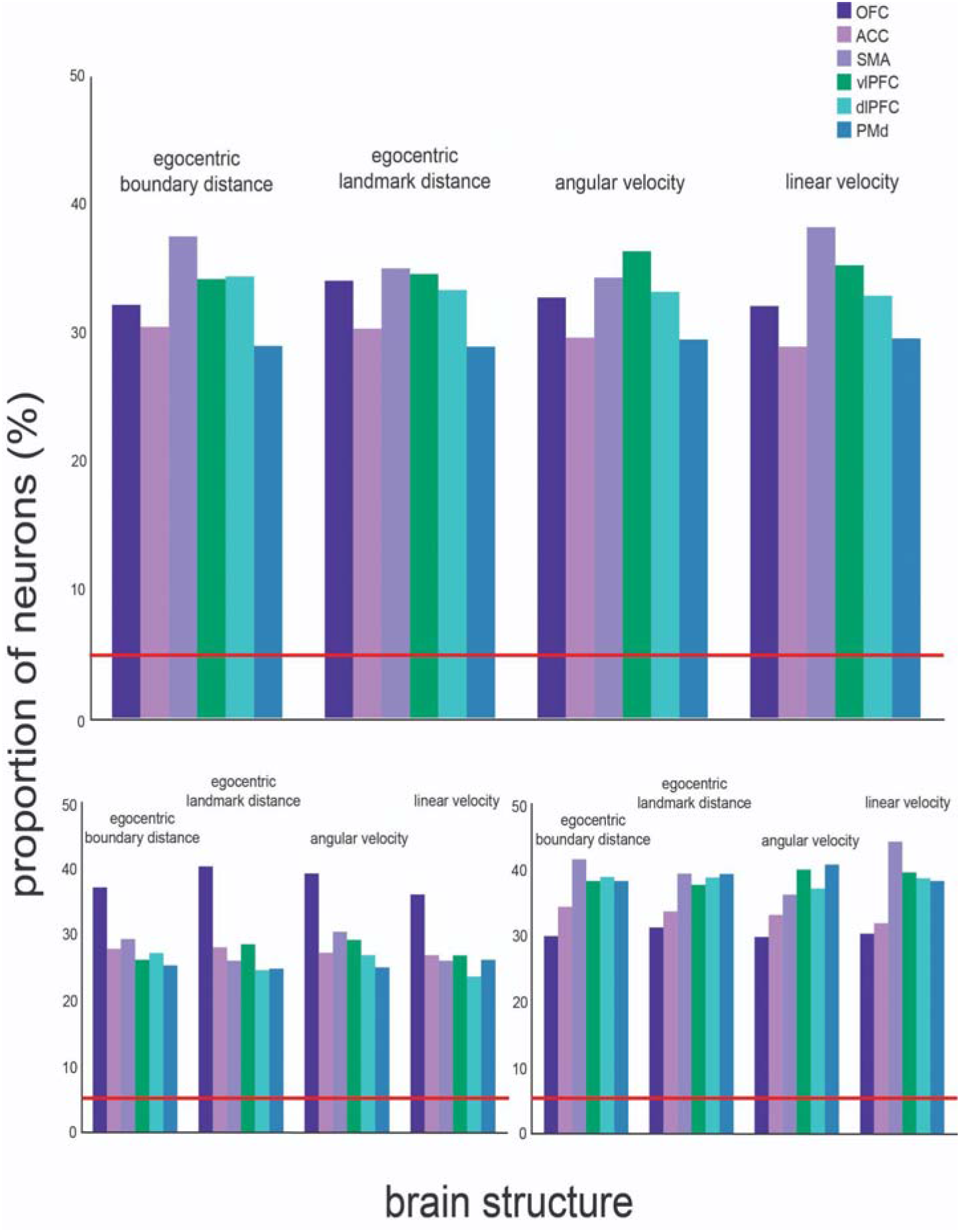
Prefrontal encoding of other navigational variables. A summary of the proportion of total recorded neurons tuned to egocentric boundary distance, egocentric landmark (feeder) distance, angular velocity, and linear velocity. Top panel: the proportions collapsed across subjects; bottom panels: the proportions within each subject. Red line: chance.

We measured linear velocity as the change in volumetric position from sample to sample (converted to cm/s). We found that a significant proportion of neurons in OFC were tuned to linear velocity (32.07%; *p* < 0.0001; binomial test). We found similar results in all 5 other areas (**Figure 4**). We found that, in all 6 structures, linear velocity tuning in the first half of the session was positively correlated with that in the second half of the session (Pearson correlation; OFC: r^2^ = 0.865; ACC: r^2^ = 0.865; SMA: r^2^ = 0.518; vlPFC: r^2^ = 0.757; dlPFC: r^2^ = 0.624; PMd: r^2^ = 0.548; *p* < 0.0001, in all cases).

To measure angular velocity, the speed with which subjects’ orientations change, we computed the change in orientation from frame to frame along both the yaw and pitch angles (converted to degrees/s). We found that a significant proportion of neurons in OFC were tuned to angular velocity (32.73%). We found similar results across all 5 other structures (*p* < 0.0001 in all cases, binomial test; **Figure 4**). As a confirmation of stability in tuning, we computed Pearson’s correlation between encoding magnitudes for each neuron to a given variable during the first and second half of the session. We found that, in all 6 structures, angular velocity tuning in the first half of the session was positively correlated with that in the second half of the session (OFC: r^2^ = 0.884; ACC: r^2^ = 0.865; SMA: r^2^ = 0.518; vlPFC: r^2^ = 0.774; dlPFC: r^2^ = 0.608; PMd: r^2^ = 0.563; *p* < 0.0001, in all cases).

Finally, we assessed selectivity for position of nearest feeder. This analysis is important because it allows us to control for possible confounder effects from reward position encoding; Egocentric landmark distance is a subject’s current volumetric distance to both the nearest and furthest feeder location in the environment. Briefly, for each timestamp, we computed the distance between the subject’s head and all four feeders. We then identified both the least and greatest distance to a feeder. We found that a significant proportion of neurons in OFC were tuned to egocentric landmark distance (34.05%; *p* < 0.0001; binomial test). We found similar results in all 5 other areas (**Figure 4**). We found that, in all 6 structures, egocentric landmark tuning in the first half of the session was positively correlated with that in the second half of the session (OFC: r^2^ = 0.846; ACC: r^2^ = 0.865; SMA: r^2^ = 0.504; vlPFC: r^2^ = 0.757; dlPFC: r^2^ = 0.593; PMd: r^2^ = 0.75; *p* < 0.0001, in all cases). The structure of our model means that even though there is coding for position to nearest feeder, the other tunings are not accounted for by this factor.

### Relationship between encoding of navigational variables and task variables

It is generally understood that neurons in these prefrontal brain regions carry information related to key behavioral variables, such as reward and choice (e.g., Kennerley & Wallis, 2009; Rushworth et al., 2011; Maisson et al., 2021). We wondered, therefore, how the robust encoding of navigational variables relates to these more familiar non-navigational task variable encodings. To this end, we computed neural encoding for five major task variables: (1) lever pressing (vs. not pressing; note that this is perfectly confounded with reward receipt), (2) number of rewards remaining throughout the entire environment, (3) number of rewards remaining at the current patch before depletion, (4) stay/leave choice after the previous trial, and (5) the predicted probability of stay/leave at each press given the number of presses remaining through the environment.

Incorporating lever pressing (i.e. the receipt of a reward) into our LN-GAM model, we found that, in OFC, 21.19% (n = 257/1213 neurons) showed selectivity to lever pressing. This proportion is significantly greater than chance (*p* < 0.0001; binomial test). We found similar and significant proportions in both subjects individually (Y: 22.55%, W: 20.69%). These proportions are roughly in line with the proportion observed in this region in other tasks (reviewed in Padoa-Schioppa, 2011). We found similar results in all 5 remaining structures (**Figure 5**).

**Figure 5.**
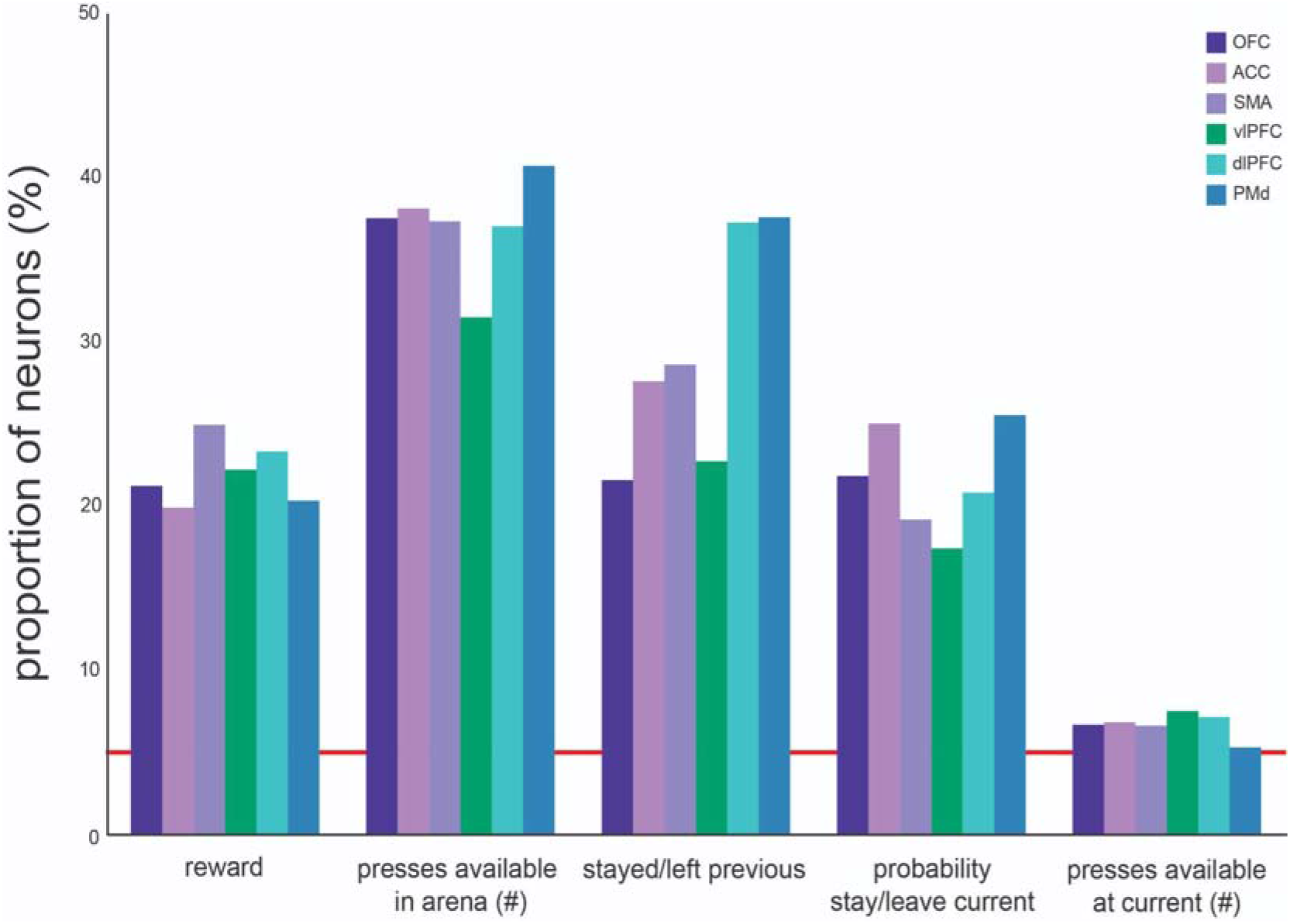
Encoding of non-navigational, task variables. Proportion of neurons significantly tuned to task-related variables. The height of the bar indicates the proportion of total recorded neurons. Each bar indicates the proportion of tuned neurons in one of the six structures. The red line indicates chance-level proportions.

To measure variables 2-5, we segmented each lever press into trial, comparable to the format expected from a chaired-task paradigm. We computed the average firing rates for each neuron per lever press and regressed the averages on the four other task variables. For each trial, we calculated how many total rewarded presses were available across all water patches in the environment. We found, for example, that a significant proportion of OFC neurons encoded the number of rewards remaining across the environment (37.59%; *p* < 0.0001, binomial test). We repeated the analysis on neural activity recorded from the other 5 areas and found similar results everywhere (**Figure 5**). For each trial, we measured the stay/leave choice from the previous trial. We found that a significant proportion of OFC neurons encoded the stay/leave choice (21.62%; *p* < 0.0001, binomial test). We again found similar results everywhere (**Figure 5**).

To measure the probability of leaving a given patch, we fitted a sigmoid function to choice by regressing the total number of presses remaining across all patches in the environment against the stay/leave decision (for details see **Methods**). We then used the fitted function to estimate the probability of staying/leaving, based on the total number of presses available in the arena. We found that a significant proportion of OFC neurons encoded the probability of leaving the current patch, while controlling for the number of rewards remaining at the current patch (21.89%; *p* < 0.0001, binomial test). We repeated the analysis on neural activity recorded from the other 5 areas and found similar results everywhere (**Figure 5**).

Finally, the number of remaining rewards at a current patch was determined as, for each press, how many presses remained before depletion at the current patch. We found that a significant proportion of OFC neurons encoded the probability of leaving the current patch, while controlling for the number of rewards remaining at the current patch (6.71%; *p* = 0.004, binomial test). We repeated the analysis on neural activity recorded from the other 5 areas and found similar results in the dACC, vlPFC, and dlPFC (**Figure 5**). The proportion of neurons encoding the number of remaining rewards at the current patch failed to reach significance in the SMA (6.64%, *p* = 0.07, binomial test) and PMd (5.33%, *p* = 0.25, binomial test).

### Variable encoding is distributed randomly among neurons within structures

Next, we asked whether navigational and task variables are encoded in distinct subpopulations of neurons. To measure the distribution of variable encoding across the recorded population within a structure, we adapted the elliptical Projection Angle Index of Response Similarity (ePAIRS) analysis (Hirokawa et al., 2019). This statistical method tells whether a group of neurons naturally falls into discrete functional subpopulations of cells. We used the LN-GAM-generated parameter weights to compute predicted variable tuning. We reduced the dimensionality and used the ePAIRS analysis to compute a cluster score (C_idx_) from the 10 best explanatory principal components (see **Methods;**Raposo et al., 2014). Given perfect clustering (C_idx_ = 1), the angle of each point is identical to its nearest neighbors. Conversely, given a perfectly random distribution of variables, we would observe a C_idx_ = 0. Indeed, we could potentially observe a C_idx_ < 0, which would indicate an even smoother distribution in variable encoding than would be generated by the data-derived null distribution (Raposo et al., 2014). This approach compares response similarity across navigational and task variables by allowing the principal component axes, rather than those of any particular variables, to determine functional clustering (Raposo et al., 2014). Clustering along PC axes prevents us from assuming any one of the measured navigational variables is necessarily independent of, or orthogonal to, another.

In the OFC, the distribution of parameter encoding was more smoothly distributed than the null distribution (score = −0.019, *p* < 0.0001; two-tailed rank sum test). This result provides strong evidence against the hypothesis that neurons are clustered into functional subtypes. Instead, it suggests that encoding any navigational variable can be achieved with any arbitrary set of available neurons. Indeed, we found similar results - significantly negative ePAIRS score - in all structures (**Figure 6A**).

**Figure 6.**
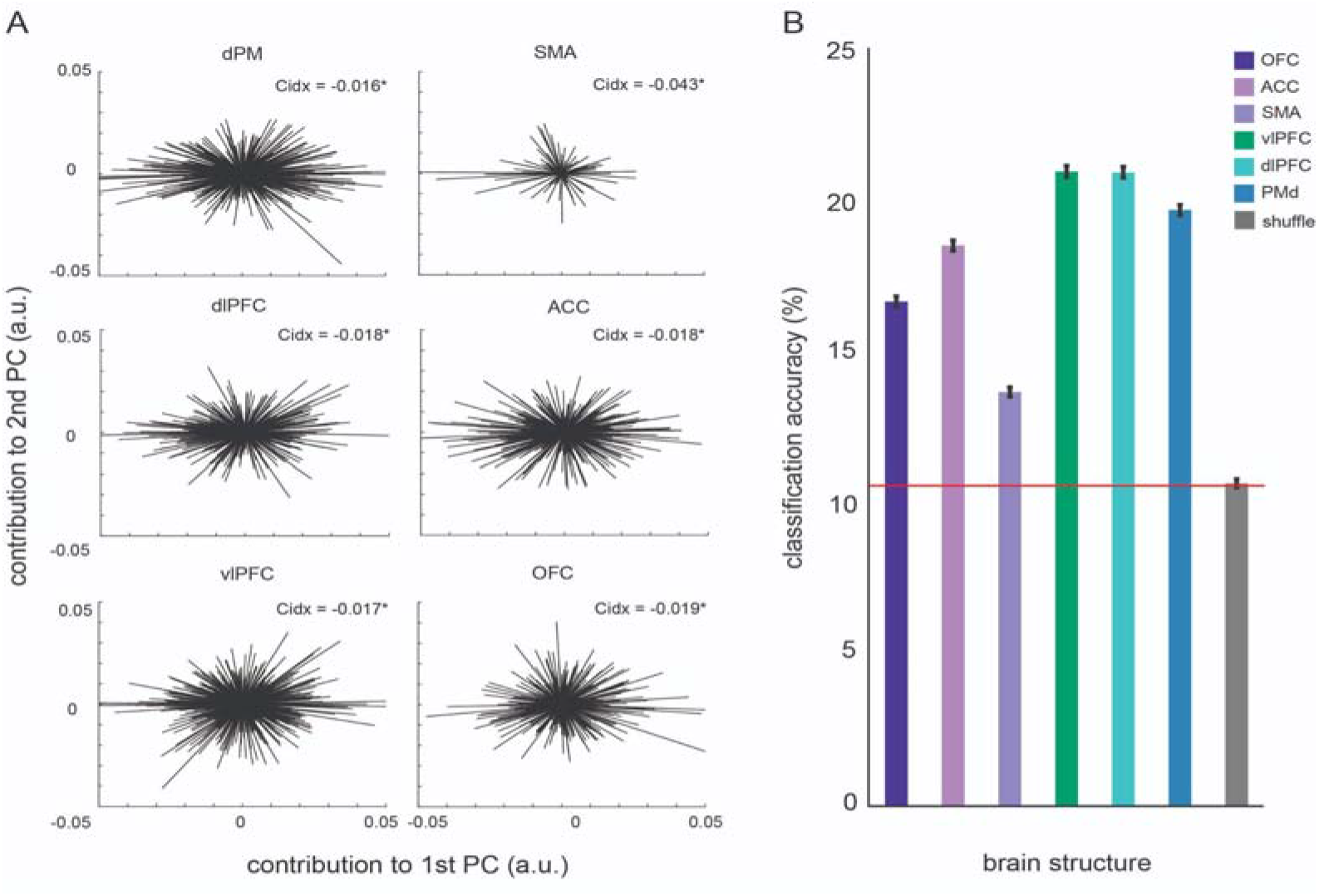
Prefrontal encoding of other navigational variables. **(A)** Weights for the first and second principal components by reducing the dimensionality of the LN-GAM derived predicted navigational variable encoding. The roughly uniform circular distributions indicate a lack of encoding-based clustering in N-dimensional principal component space. The inlaid score indicated the clustering index from the ePAIRS analysis and an asterisk denotes significance at *p* < 0.05 **(B)** Accuracy of a linear decoder, a regression-based support vector machine, in using population neural activity to decode the 2D position of the subject. Each bar indicates the accuracy for decoding subject position based on neural activity in each structure. Gray: shuffled values. Bars: standard error across bootstrapped iterations. Red line: chance.

Why might the brain use such strongly multiplexed encoding? One advantage is that it allows the same population of neurons to meet distinct behavioral demands, while continuing to support simple linear combinations for downstream decoding (Raposo et al., 2014; Fusi et al., 2016). In one study, authors demonstrated that, despite the category-free coding they observed in the posterior parietal cortex, it was possible to nonetheless perform a linear readout of firing to estimate task variables, demonstrating that category-free coding does not result in a sacrifice in fidelity (Raposo et al., 2014). To confirm that the same applies to our own data, we performed the same control analysis described in that study. To do so, we asked whether we could use a linear decoder to accurately predict the position of the animal, based on the firing rates across the population (see **Methods**). We segmented the arena into 9 decodable zones, so chance-level decoding accuracy was 11.11% (i.e., 1/9). We compared the decoding accuracy from population activity in each structure with the chance level, after correcting for multiple comparisons (that is, we used ⍰ = 0.0083). We found that position could be reliably decoded from population neural activity in the OFC (*M* = 16.64%, *SEM* = 0.17%; *p* < 0.001, two-sample t test). We found similar results in all structures (**Figure 6B**). These results indicate that the random distribution of encoded parameters allows for simple linear decoding, likely by allowing for the arbitrary recruitment of available neurons to combine parameters and compute navigational variables. Note that all of these analyses were repeated and confirmed for each subject individually (data not shown).

### Coding of all measured variables grows along a ventral to dorsal gradient

We have previously shown that encoding of several economic variables exhibits a characteristic functional ventral-to-dorsal gradient, in which choice-relevant information increases in amplitude as it moves dorsally along the medial wall of the PFC (Maisson et al., 2021). We next asked whether such a functional gradient is also observed for navigational variables. We identified two distinct potential ventral-to-dorsal anatomical gradients in our data: a medial series (OFC → dACC → SMA → PMd) and a lateral series (OFC → vlPFC → dlPFC → PMd).

We found that the strength of position encoding increases along both gradients. Formally, the regression weight of response strength, computed for each neuron, increased against a dummy variable for the structure’s hierarchical position (medial series: ρ^2^ = 0.067; lateral series: ρ^2^ = 0.051; *p* < 0.0001 in both cases, Spearman’s correlation). Note that we observed the same pattern in an analysis that ignored brain area and used the depth of the electrode (medial series: r^2^ = 0.055; lateral series: r^2^ = 0.065; *p* < 0.0001 in both cases, Pearson’s correlation). This second result indicates that the particular boundaries we chose for brain area were not the determining factor for establishing our measured gradients. In other words, regardless of the particular parcellation one chooses, the more dorsally located either the brain area or neuron, the stronger the position encoding.

We also found a similar functional gradient for head direction encoding (medial series: ρ^2^ = 0.099; lateral series: ρ^2^ = 0.078; *p* < 0.0001, Spearman’s correlation) and observed the same pattern in an analysis that ignored brain area and used the depth of the electrode (medial series: r^2^ = 0.081; lateral series: r^2^ = 0.095; *p* < 0.0001 in both cases). Likewise, we found similar results for egocentric boundary distance tuning (medial series: ρ^2^ = 0.125; lateral series: ρ^2^ = 0.099; *p* < 0.0001) and in an analysis that used the depth of the electrode (medial series: r^2^ = 0.098; lateral series: r^2^ = 0.076; *p* < 0.0001 in both cases). We found similar results for angular velocity (medial series: ρ^2^ = 0.125; lateral series: ρ^2^ = 0.099; *p* < 0.0001), and in an analysis that used the depth of the electrode (medial series: r^2^ = 0.098; lateral series: r^2^ = 0.076; *p* < 0.0001 in both cases). We also found a similar functional gradient for linear velocity (medial series: ρ^2^ = 0.118; lateral series: ρ^2^ = 0.099; *p* < 0.0001), and in analysis that used the depth of the electrode (medial series: r^2^ = 0.097; lateral series: r^2^ = 0.071; *p* < 0.0001 in both cases).

We also found similar patterns for our non-navigational task variables. We found that the encoding magnitude of stay/leave choice probability slightly but significantly increased along both the medial and lateral wall gradients (medial series: ρ^2^ = 0.071; lateral series: ρ^2^ = 0.069; *p* < 0.0001, Spearman’s correlation). We found that the unsigned weight of reward encoding, as extracted from the LN-GAM approach, increased along both the medial and lateral wall gradients (medial series: ρ^2^ = 0.081; lateral series: ρ^2^ = 0.067; *p* < 0.0001, Spearman’s correlation). As before, these same patterns were significant when using electrode depth (*p* < 0.0001 in all cases, Pearson’s correlation).

## DISCUSSION

We investigated the neuronal encoding of navigational variables while macaques moved freely in a large environment. We find that single neurons in six prefrontal areas robustly encode several navigational variables. These results qualitatively resemble those obtained in a previous macaque study using similar analysis methods but focusing on the hippocampus (Mao et al., 2021). Moreover, our analyses indicate that navigational and non-navigational variables are encoded in the same set of neurons, using principles of mixed selectivity (Raposo et al., 2014; Fusi et al., 2016; Blanchard et al., 2018). The widespread nature of encoding of navigational variables outside of the hippocampal complex suggests that processing of navigational information may not be narrowly localized to a small portion of the brain, but instead may be a more general property of the brain, including the frontal lobes. More generally, these results tie into emerging theories that see navigation as, essentially, a special case of associative mapping (Behrens et al., 2018; Epstein et al., 2017; Whittington et al., 2020; Stachenfeld et al., 2017; Schuck and Niv, 2019). From this perspective, navigational information may be found in prefrontal regions because of their more general role as flexible encoders of associative information.

Several recent reports in rodents have identified encoding of navigational variables beyond the hippocampal complex (Basu et al., 2021; Long and Zhang, 2021; Poo et al., 2022; Wikenheiser et al., 2021). Likewise, a small but growing literature in humans emphasizes the potential roles of non-hippocampal regions, especially prefrontal ones, in navigation (reviewed in Patai and Spiers, 2021). Our report therefore adds to this literature. Some of these results have proposed that navigational information outside of hippocampus may play a distinct role, such as linking space to value or changing plans; however, existing data are also consistent with the hypothesis that hippocampal and prefrontal regions play largely overlapping roles. Future work will be needed to disambiguate these hypotheses.

We find that encoding of navigational variables coexists with encoding of more often studied non-navigational task variables, indeed, even in the same neurons. These results are important because they indicate that navigational encoding is not part of specialized patches or neural subregions, nor of distinct neurons that were inaccessible in prior studies. Instead, the information about the navigational world is fully intertwined with information about other aspects of the task.

The ventral-to-dorsal gradient in coding strength we report here resembles one we have found, in a narrower set of analyses on non-navigational variables, for the medial wall series (Maisson et al., 2021). Why would the brain have stronger encoding more dorsally? We propose that the information is likely to be present at all levels but is encoded in a format that is more accessible to the motor system (and likewise to our decoding analyses) in more ventral structures. In other words, more dorsal structures show information relevant to action in a more untangled manner (Yoo & Hayden, 2018). Our results here support this proposal and show that it applies to both the medial series (as we previously showed) and to the lateral series (which we did not). Conversely, these data argue against a functionally modular arrangement, in which each area has a specific nameable function, such as “evaluate,” “compare,” and “control”. At the same time, these data argue against a purely distributed (or Lashleyan) view, in which all information is present in the same form at all times across prefrontal cortex. Instead, they support a third, hierarchical view (Fuster, 2000 and 2001; Hunt et al., 2014 and 2018; Hunt and Hayden, 2017; Rushworth et al., 2011).

One major limitation of the present study is that we did not measure gaze direction. As such, we could not control for variability in gaze direction as a possible confounder. In a recent, important paper, Mao et al. (2019) characterized navigational tuning in the hippocampus of freely moving primates. They showed the results of both a traditional tuning curve and the LN-GAM approach used here. The traditional tuning curve approach, using a single fitted variable at a time, yielded tuned proportion analogous to those commonly reported in the rodent HC. Critically, however, the LN-GAM approach included a variable for “facing location” within the large, simultaneously fitted model. They demonstrated that, while neurons continued to be selective to other navigational variables, some hippocampal tuning was preferentially selective to facing location (see also Killian et al., 2012 and Jacobs et al., 2013). In future studies we hope to investigate the relationship between location and viewed location.

Ultimately, we believe that these results provide a strong argument for the use of naturalistic tasks. It is well known that natural behavior is continuous, complex (multi-effector), embedded (in an environment), and seldom isolated from other competing demands on attention and planning (Brown and De Bivort, 2018; Datta et al., 2019; Krakauer et al., 2017; Matusz et al., 2019; Pearson et al., 2014; Yoo et al., 2021). Nonetheless, most laboratory tasks, including the great bulk of our own past work, does not hew to these principles. Instead they use overly simplified laboratory tasks. Simple laboratory tasks have great benefits, especially in tractability, but they have disadvantages. One is that they often do not manipulate variables that may, if manipulated, be found to be major drivers of neural activity. Ignoring that tuning may in turn obscure the broader and more general functions of neurons in regions of interest.

## METHODS

### Surgical procedures

The University Committee on Animal Resources at the University of Minnesota approved all animal procedures. Animal procedures were designed and conducted in compliance with the Public Health Service’s Guide for the Care and Use of Animals and approved by the institutional animal care and use committee (IACUC) of the University of Minnesota. Two male rhesus macaques (*Macaca mulatta*) served as subjects. Animals were habituated to laboratory conditions, trained to enter and exit the open arena, and then trained to operate the water dispensers. We placed a cranium adherent form-fitted Gray Matter (Gray Matter Research) recording chamber and a 128-channel microdrive recording system over the area of interest. We verified positioning by reconciling preoperative magnetic resonance imaging (MRI) as well as naive skull computed tomography images (CT) with postoperative CTs. Animals received appropriate analgesics and antibiotics after all procedures.

### Recording sites

We approached our brain regions by controlled and monitored advancement of individual electrodes, through reconciliation of measured thread-count against postoperative CT images. The implanted microdrive was connected through jumper cables and a head stage to a removable data logger (SpikeGadgets). The data logger was wirelessly triggered to store neural recordings onto a removable memory card from which it was then extracted at the end of each session.

We defined **OFC** as lying within the coronal planes situated between 39.6 and 23.9 mm rostral to the central sulcus, the horizontal planes situated between 23.3 and 45.7 mm from the brain’s dorsal surface, and the sagittal planes between 1.3 and 22.1 mm from the medial wall.

We defined **dACC** as lying within the coronal planes situated between 33.5 and 7.4 mm rostral to the central sulcus, the horizontal planes situated between 11.1 and 38.5 mm from the brain’s dorsal surface, and the sagittal planes between 0.4 and 6.7 mm from the medial wall.

We defined **SMA** as lying within the coronal planes situated between 30.4 and 13.1 mm rostral to the central sulcus, the horizontal planes situated between 16.2 and 34.3 mm from the brain’s dorsal surface, and the sagittal planes between 0.4 and 6.7 mm from the medial wall.

We defined **vlPFC** as lying within the coronal planes situated between 46.8 and 16.8 mm rostral to the central sulcus, the horizontal planes situated between 16.2 and 47.2 mm from the brain’s dorsal surface, and the sagittal planes between 1.1 and 19.4 mm from the medial wall.

We defined **dlPFC** as lying within the coronal planes situated between 36.4 and 12.3 mm rostral to the central sulcus, the horizontal planes situated between 17.4 and 48.9 mm from the brain’s dorsal surface, and the sagittal planes between 1.1 and 19.4 mm from the medial wall.

We defined **PMd** as lying within the coronal planes situated between 26.7 and 2.9 mm rostral to the central sulcus, the horizontal planes situated between 6.8 and 37.9 mm from the brain’s dorsal surface, and the sagittal planes between 1.9 and 23.7 mm from the medial wall.

### Behavioral task

There were four statically positioned water dispensers (“patches”) available with programmed delivery schedules. Each patch delivered a fixed amount of 1.5mL per lever press. The first four presses, regardless of patch sequencing, were rewarded with water delivery. The fifth lever press was unrewarded and led to a 3-minute deactivation of the patch. That is, animals could press fewer than five times, leave to engage with a different patch, and return to the same patch in the state they left it. No reset or deactivation was applied if the animal left the patch. A patch was only reset if the subject pressed the lever a fifth time and waited 3 minutes for it to reactivate. Each rewarded press followed the same programmed sequence. An activate patch was indicated by a fully blue display. A lever press changed the display to white with a green plus-sign in the center, an auditory cue was played, and a solenoid opened to dispense reward. After dispensing the solenoid closed, the auditory cue ended, and the green plus-sign disappeared. The screen remained white for 2 additional seconds before the screen turned blue again to indicate the availability of another lever press. The fifth lever press was instead followed by the screen immediately turning white, with no visual or auditory reward cue and no water delivery. However, it is important to note that task engagement was not required; a subject could choose not to engage with the patches for the entirety of the session. Otherwise, the measured behavior was simply the free movement of the subject through the arena. On average, subjects pressed levers 136 +/− 8 times (subject Y: 166 +/− 7, subject W: 107 +/− 9), amounting to roughly 204 mL of water, per session, received for interacting with patches. Prior and concurrent chaired task training of these subjects included two risky choice tasks (Azab and Hayden, 2018; Farashahi et al., 2019; Yoo et al., 2020; Wang et al., 2019 and 2022) and a simpler choice task (Blanchard et al., 2014; Ebitz et al., 2019).

### Behavioral analysis

There were 5-7 patch events recorded for each patch engagement: lever press, screen off (set to white), screen/auditory reward cue (rewarded presses only), solenoid open (rewarded presses only), solenoid close (rewarded presses only), screen off (set to white), timeout (deactivation press only), and screen reset. Lever press events that led to a reward were used to analyze the reward encoding. To track the position of the animal, we employed our custom integrated software/hardware markerless tracking and pose estimation system, the OpenMonkeyStudio (Bala et al., 2020; Hayden et al., 2021).

To measure movement through space we extracted the volumetric position of the head during each session. Each of the axes (X, Y, and Z) were extracted as independent one-dimensional parameters. The X and Y axes were aggregated into a *2D position* parameter, and the Z axis was used to measure *head elevation*. Together, these 2 parameters defined *3D position*. To measure the 3 angular axes of the head, we first centered the extracted nose and neck coordinates, placing the estimated head position at the volumetric axis-origin. For each frame, we applied Euler angle transformations on the head-fixed coplanar points to determine the yaw, pitch and roll angles between the origin and the current head orientation. We defined the *head direction* parameter as a 1D vector for the yaw angle of the head. We defined the *head tilt* parameter as a 2D vector consisting of the pitch and roll angles. We measured egocentric boundary in polar coordinates, defined as a 2D parameter consisting of the radial distance between the subject and the nearest wall on the azimuthal plane and the angle of the subject relative to the center of the room. To measure angular velocity, we computed a two-dimensional parameter, where the first dimension reflects the rotation speed of the head (degrees/second) along the yaw angle and the second dimension reflects the rotation along the pitch angle. To measure linear velocity, we computed the distance traveled (centimeters/second) in any direction.

### Linear-Nonlinear Poisson-distributed Generalized Additive Model

We adapted an approach previously developed for estimating the simultaneous tuning of neural activity to multiple binary, linear, and radial parameters (Hardcastle et al., 2017; Laurens et al., 2019; Balzani et al., 2020; Mao et al., 2021; Ledergerber et al., 2021; Yoo et al., 2020 and 2021). Briefly, single unit recordings and extracted parameters, from 100-min sessions, were binned into 100-ms time bins. Neural data were fitted to multiple, nested linear-nonlinear-Poisson distributed generalized additive models. Data were divided into 10 chunks across the session. Within each chunk, model estimates were computed using 5-fold cross-validation, by dividing each chunk into 5 more subchunks. A given subchunk from each chunk was concatenated to constitute the test set (20% of the data) and the remaining subchunks were concatenated to constitute the training set (80% of the data). The process was repeated for each given subchunk. The estimated log-likelihood quantified the model performance, and the distribution of log-likelihood estimates was compared to a null distribution (one-sided Wilcoxon signed rank test, alpha = 0.05).

The best-fitting model was selected using an optimized forward search. Models containing only one parameter (1^st^ order models) were fitted first. If these models performed better than the null, models containing two parameters (2^nd^ order models) were fitted. The process was repeated until the performance of the newly fitted model no longer improved over the previously fitted model. A neuron was categorized as significantly tuned to a given parameter, if that parameter was included within the best-fitting model. The proportion of neurons encoding each parameter was calculated from the number of neurons whose best-fitting model included the given parameter. Importantly, the additive process for multiple terms was incorporated within the exponentiation used to compute the estimated firing, thus making selectivity to multiple parameters a conjunctive (nonlinear) estimate.

We validated the model on our data through a series of controls. After fitting the LN-GAM models, we randomly selected a neuron that was not significantly tuned to any combination of parameters. We modified the spike train for that neuron to include a single spurious high firing event, once in each of 3 time periods. We also selected both 10 and 100 bins from each of 3 time periods to insert spurious high firing. Finally, we time-locked the insertion of 1, 10, and 100 simulated spurious events to consistent instances of the tracked position. We repeated this process across 20 randomly selected, untuned neurons from each structure, generating a set of 1200 synthetic datasets. We fitted each of these simulated time series against the original parameters. We then determined the proportion of fitting processes that resulted in significant tuning. Only 8.08% (n = 97/1200) of control sets generated a statistically significant fit. These controls confirmed that the best-fitting models were not being influenced by random or spurious events, and instead reflected reliable tuning to the selected parameters.

### Stay/Leave choice probability and neural encoding

We adapted an approach commonly used in traditional neuroeconomics paradigms (e.g., Strait et al., 2014; Maisson et al., 2021; Maisson et al., 2022). We leveraged the strictly timed structure of the foraging task to isolate the 2-second epoch during which subjects were rewarded for pressing a lever. Each lever press is necessarily separated from the next by a minimum of 4 seconds; 2 seconds of a reward period and 2 seconds of an intertrial interval. Therefore, each rewarded lever press was treated as a separate trial. We aggregated all trials for each feeder across the session. For each trial, we determined the number of rewarded presses remaining at both the current patch and all distal patches. We also determined whether the current trial occurred at the same or different patch from the previous trial. If it was the same as the previous patch, the subject was said to have made a “stay” choice on the previous trial. Conversely, if it was at a different patch, the subject was said to have made a “leave” choice on the previous trial. This binary choice variable was fitted to the total number of rewarded presses available across the distal patches, using a logistic regression. The resulting sigmoidal function was then used to estimate the stay/leave probability for each trial, based on how many rewarded presses were available elsewhere in the arena. Finally, for each neuron and each trial, an average firing rate was computed from the same post-press epoch. The trial-length vector of stay/leave probability was used to predict this trial-length vector of firing rates, using a linear regression that simultaneously controlled for the number of rewarded presses remaining at the current patch.

### Distribution of encoding parameters

We adapted an approach previously developed for estimating clustering in n-dimensions, the elliptical Projection Angle Index of Response Similarity (ePAIRS; Raposo et al., 2014; Hirokawa et al., 2019). Briefly, the activity of a neuron in any given session may show preferential tuning to a particular parameter, or condition. If neurons form specialized subpopulations that are reliably tuned to common variables, then they should form clusters along a shared axis that consistently describe their variance. Thus, reducing the dimensionality of the tuning conditions across the population code should elucidate a functional cluster. That is, where an animal is located in X-Y space may change from day to day, thus changing the shape of neuronal tuning on that day. Yet, a specialized “position-tuned” subpopulation should share a common “position axis” that explains the variance in their activity.

To define a tuning condition, we compute the estimated marginal means (EMMs) from the full-fitted LN-GAM model. Estimates were computed using the minimum and maximum values for each parameter in the design matrix. We decided to use the fully-fitted model estimates because it reduces sparsity in the population matrix and should reflect the relative weights of the significantly predictive parameters. We reduce the dimensionality of the tuned conditions using principal component analysis (PCA) from across the recorded population and computed the best 10 PC loadings for each neuron. We projected the loadings onto the surface of an n-dimensional ellipsoid, computed the relative angles of each point, and compared the angular distance between all pairs of nearest neighbors. A functional cluster, therefore, should have relatively tight angles between nearest neighbors when compared to a random distribution. Statistical significance was computed using a rank sum test to compare the distribution of nearest neighbor angles with the random distribution.

In each session, a given neuron might be more heavily tuned to one parameter than another. That is, the EMMs reflect the distribution of preferred parameters. We would expect these estimates to show clustering along the dimensions of the preferred parameters. Therefore, as a positive control we performed the ePAIRS analysis on the EMMs and found significant clustering in all areas (*p* < 0.001, in all areas; two-tailed rank sum test), confirming that clustering would be detectable in our data if it were present.

### Linear decoder

To determine if randomly distributed parameter encoding continues to support downstream decodability from linear parameter combinations, we implemented a regression-based support vector machine. Tracking data from each session was spatially binned into 9 continuous zones. We define a trial as an entry into the zone, followed by continuous zone occupancy until exiting the zone. Timestamps in the neural activity from a given session were locked to these occupancy-based trials and separated accordingly. Thus, each session consisted of zoned trials for which peristimulus time histograms could be computed. The average firing rate across each trial was computed independently for the trials corresponding to the occupancy in each zone. We constructed a pseudopopulation of pseudotrials, 495-trials X n-neurons, by randomly selecting 55 trials from each zone for each neuron. We repeated the random bootstrapping procedure twice to constitute one training set and one test set. We created two more matrices by shuffling the zone labels. A regression-based SVM model was trained on the training set and predictions were calculated by fitting the model to the test set. Accuracy was measured as the overall rate of successfully predicting the zone from the population neural activity in the test set. We repeated this process 500 times and averaged the classification accuracy across iterations. We then compared this average accuracy to that from the shuffled data.

### Gaussian Mixture Models, Expectation-Maximization, and clustering

We developed, to fit our neural data and behavioral variables, a machine learning approach to compare data-driven clustering to supervised-clustering. In this case, the supervised-clustering approach was to simply group all the navigational parameter encoding weights by the known gross anatomical structure for the respective neuron. In the unsupervised approach, we fit a set of Gaussian Mixture Models (GMM), optimized them, and used them to determine clusters. We started with two proofs of concept. In the first, we demonstrated that fundamentally distinct groups in the data formed fundamentally separable clusters. To do this, we generated and aggregated 6 independent distributions, each 1000 (samples) by 11 (variables). Each distribution had randomly defined means and covariance structures for each variable. We decided on 6 because we knew we were going to compare the results to the *a priori* area-based clusters. We then performed a principal component analysis and projected the data into principal component space. A GMM was fitted to this artificial data using an Expectation-Maximization (E-M) algorithm. The E-M algorithm was initiated with a randomly selected seed and given 1000 attempts to converge at a minimum.

To reduce the risk of converging to a local minimum, this process was itself repeated 1000 times before the final model was selected. This process was repeated for each number of possible clusters (from 1 to 6). The best fitting model, and thus how many components it defined, was selected based on which model offered the lowest AIC and BIC scores. Finally, the best model was used to cluster the data. The cluster index was then compared to the known *a priori* group label (i.e. the brain area in the case of real data), and the alignment between cluster and group label was determined as the rate of match. This process was then repeated over 1000 iterations, and the mean and standard error were computed across iterations. This approach was shown to generate perfect alignment, as expected. To determine the minimal possible alignment, we followed the exact same procedure, except that we forced the mean and covariance structures of the 6 initial distributions to be identical. As expected, this invariably generates a single cluster and thus offers the minimal possible alignment.

We argued that if brain areas defined meaningful functional boundaries, then the distributions of variable encoding weights should be distinct for each brain area. To establish a control group, we generated a synthetic data set using the mean and covariance structures of our real data but with random points along that distribution structure. This synthetic data should be fitted by GMMs that have the same number of components (6) as the brain areas they come from, and the assigned cluster label should align fairly closely with the *a priori* brain area label. We repeated the exact same process as described above using these synthetic data sets. Finally, to get the true clustering, we performed the exact same procedures but fitting the GMMs to the real distributions.

## Acknowledgements

We thank the Hayden/Zimmermann lab for valuable discussions as well as Brenna Knaebe for help with animal care and preparation. This work was supported by NIH grants R01 MH128177 (JZ), P30 DA048742 (JZ, BH), R01 MH125377 (BH), NSF 2024581 (JZ, BH) and a UMN AIRP award (JZ, BH) from the Digital Technologies Initiative (JZ), from the Minnesota Institute of Robotics (JZ).

